# Early dynamics of excitation and inhibition maintain late frequency tuning in auditory cortex

**DOI:** 10.1101/2024.01.26.577483

**Authors:** Ashlan P. Reid, Anthony M. Zador, Tomáš Hromádka

## Abstract

In the auditory cortex the onset of a tone evokes time-varying excitation and inhibition. However, the role of early inhibition in shaping the temporal properties of tone-evoked responses has not been fully characterized. By using Archaerhodopsin-3 (Arch) to photo-suppress the activity of the parvalbumin-expressing (PV) class of inhibitory interneurons, we manipulated the early component of tone-evoked inhibition. We find that early inhibition directly controls the output gain of the response, reducing the number of spikes proportionately across all frequencies. However, by controlling early activity, transient inhibition prevents late excitation and spiking for non-optimal frequencies. Thus, transient tone-evoked inhibition plays a critical long-lasting role in shaping response properties in the auditory cortex.

## Introduction

The stimulus selectivity of a cortical neuron is determined by the interaction of excitatory and inhibitory synaptic inputs it receives. The simplest form of interaction is additive, with the output of the neuron proportional to the difference between excitation and inhibition. Such additive interactions form the basis of many artificial neural network models (Holt & Koch, 1997; McCulloch & Pitts, 1948; Silver, 2010). Other models of the neuronal transfer function include a divisive role for inhibition. The operation of division provides a neuron with many capabilities, including improving the dynamic range of the input-output transformation (Blomfield, 1974; Mitchell & Silver, 2003). These models typically assume static inputs and use time-averaged spike rates as a measure of neuronal output.

However, excitation and inhibition often vary with time. This is particularly true for synaptic events set in motion by transient sensory events. Time-varying excitatory and inhibitory inputs can interact in complex ways (Ferster & Miller, 2000; Higley & Contreras, 2006; Isaacson & Scanziani, 2011; Oswald et al., 2006; Priebe & Ferster, 2008; Stiebler et al., 1997; Swadlow & Gusev, 2000). For example, the relative timing of excitation and inhibition can be critical in shaping the sensory response, as has been proposed to underlie motion selectivity in the retina (Koch et al., 1983) and direction selectivity in the rodent barrel cortex (Wilent & Contreras, 2005). Such temporally dynamic interactions are not readily described in terms of steady state addition or division.

In the auditory cortex, the onset of a tone elicits a stereotyped barrage of excitation followed within a few milliseconds by a barrage of inhibition (Wehr & Zador, 2003; Wu et al., 2008; Zhang et al., 2003; Zhou et al., 2014). The short period before the onset of inhibition presents a “window of opportunity” during which unchecked excitation can elicit spikes. The time course of this excitation-inhibition sequence is independent of frequency, so the duration of the window should not contribute to the initial frequency tuning of the neuron. Instead, the fast tone-evoked inhibitory conductance in this system is predicted to act like a delayed “off” switch, quenching the excitatory conductance and thereby limiting the number of spikes elicited by a stimulus.

Here, we tested the consequences of eliminating early tone-evoked inhibition on the output of auditory cortical neurons. Our strategy was to selectively and reversibly silence the activity of one subclass of interneurons, defined by the expression of parvalbumin (PV), by photo-stimulation of genetically targeted Archaerhodopsin-3 (Arch) (Chow et al., 2010). We hypothesized that by controlling the activity of PV interneurons we could selectively manipulate fast tone-evoked inhibition. Evidence for the action of PV interneurons in early stimulus-evoked inhibition comes from many sensory cortices: they constitute about half of all cortical interneurons, provide powerful somatic inhibition to excitatory neurons, receive strong direct thalamic inputs, and respond with high reliability to thalamic activation (Celio, 1986; Cruikshank et al., 2007; Gibson et al., 1999; Hattori et al., 2017; Inoue & Imoto, 2006; Kerlin et al., 2010; Porter et al., 2001; Runyan et al., 2010; Schiff & Reyes, 2012; Sun et al., 2006; Wood et al., 2017).

We find that, whereas suppression of early inhibition has no effect on the initial tuning width of the tone-evoked response, it unexpectedly increases the tuning width late in the response. This delayed change is associated with an excess excitatory drive rather than a direct effect of suppressing inhibition. Thus, early inhibition contributes indirectly to the shape of tone frequency tuning in the auditory cortex over long timescales. Our results show that by enacting transient gain modulation, inhibition can influence the shape of cortical responses for tens to hundreds of milliseconds after tone response onset.

## Results

We set out to determine the role of early tone-evoked inhibition in the frequency tuning of auditory cortical neurons by selectively controlling the activity of PV interneurons. We first performed cell-attached recordings *in vivo* to examine the effects of selective and reversible optogenetic silencing of PV interneurons on tone-evoked spiking responses in the auditory cortex. We then reversibly silenced PV neurons during thalamic activation *in vitro*, to assess their role in thalamically-evoked feedforward inhibition. Finally, we reversibly silenced PV interneurons and performed whole-cell patch recordings *in vivo* to reveal the synaptic events underlying the changes observed in tone-evoked spike responses.

### Spontaneous and tone evoked spike rates increase when PV interneurons are silenced

We first studied the role of PV-mediated inhibition in shaping tone-evoked spike response properties. We used cell-attached patch recordings in anesthetized mice to record the spiking activity of single neurons in auditory cortex under interleaved control and during silencing of PV interneurons (control and PVoff conditions, Fig. 1b). Photo-suppression of PV interneurons induced a robust increase in the spontaneous and tone-evoked spike counts (Fig. 1c). Both spontaneous and tone-evoked firing rates increased significantly across the population (Fig. 1d,e; n=19 neurons, paired t-test), indicating lower inhibitory activity during the entire light pulse (Fig. 1d inset). Higher firing rates led to a significantly higher fraction of tone-responsive trials (Fig. 1f, paired t-test). By contrast, the timing of onset responses, assessed by either the median first spike latency or its standard deviation across responsive tones (see Materials and Methods), showed no change during PVoff conditions (Fig. 1g), which is consistent with the onset of responses being primarily determined by the initial arrival of excitatory activity.

**Figure 1.**
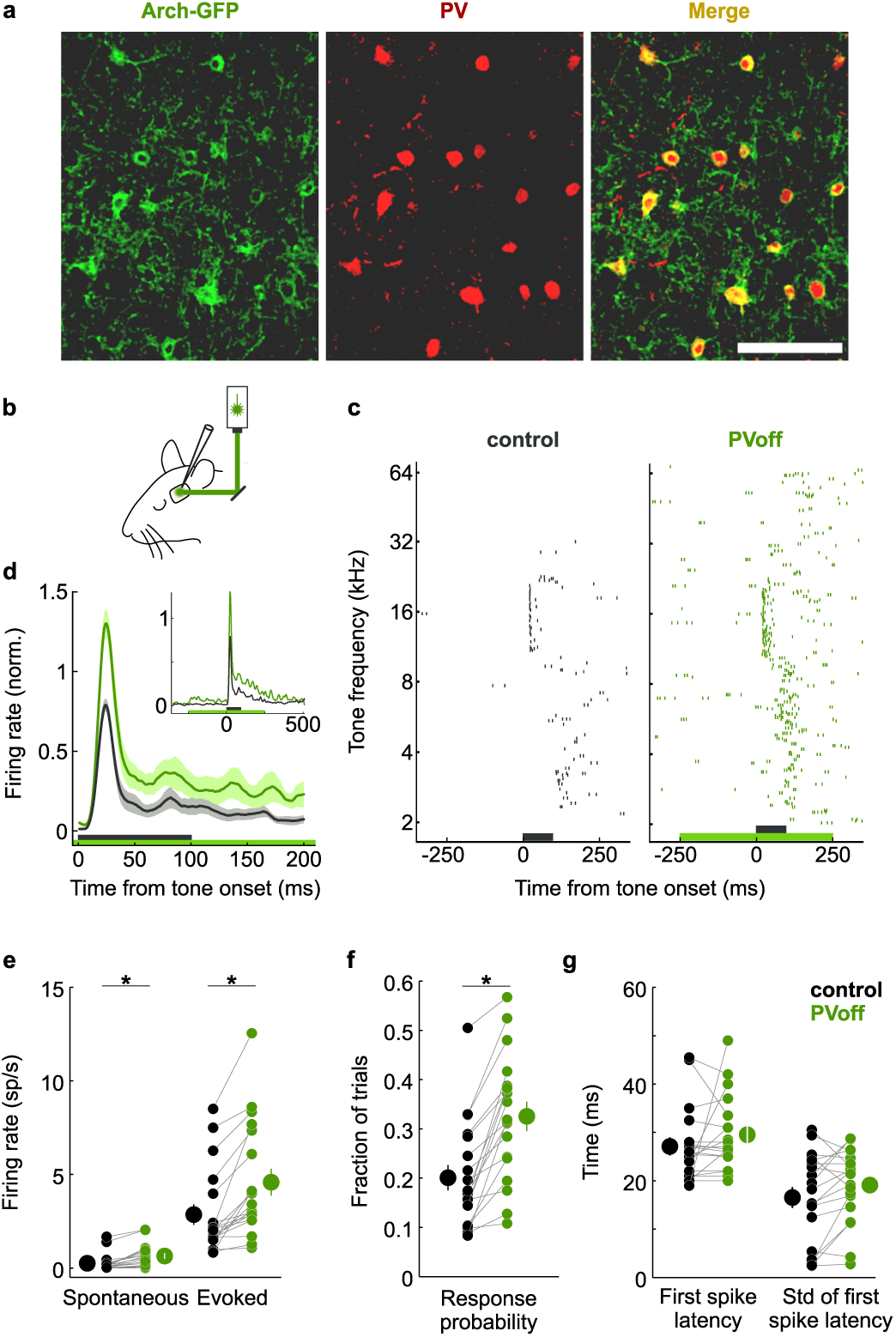
Silencing PV interneurons increased firing rates but did not change tone-evoked first spike timing. **a.** lnterneurons expressing membrane bound Arch-GFP fusion protein (green, left) were immunopositive for parvalbumin (red, middle; red/green merge, right). Scale bar indicates 100 µm. **b.** In vivo preparation showing configuration of laser light stimulation (532 nm laser, 1 mm beam width, 18-30 mW total power) and recording electrode. **c.** Silencing PV interneurons increased the number of evoked spikes. Rasters represent spikes from twenty responses elicited by 2 kHz to 64 kHz tones each presented under control (black) and test (PVoff, green) conditions. Black bar indicates tone presentation, green bar indicates light presentation. **d.** Photo-suppression of PV interneurons increased firing rates of tone-evoked responses. Average peristimulus time histogram (PSTH, n=19 neurons) computed from the spiking responses to tones (as in A) under control (black) and PVoff (green) conditions. Shaded areas show s.e.m. Black bar indicates tone presentation, green bar indicates light presentation. Inset shows the same PSTH for the full trial duration. **e.** Spontaneous (left) and evoked (right) firing rates increased significantly when PV interneurons were silenced (p<0.05, paired t-test). **f.** Fraction of trials with at least one spike during tone presentation (response probability) increased significantly under PVoff condition (p<0.05, paired t-test). **g.** Median first spike latency (left) and standard deviation of first spike latencies (right) did not change significantly under PVoff conditions (paired t-test).

We initiated each light pulse well in advance of the tone onset to separate the tone response from the transient synchronous rise in spontaneous spike rate across the cortex caused by light onset (Fig. 1d inset). As the light pulse persisted, the rate of spontaneous spikes decayed to a baseline higher than the spontaneous rate under control conditions. In control trials (500 ms light pulse without a tone), interleaved with experimental trials, there was no indication of accommodation of PV interneuron silencing over the duration of the light pulse, and further control *in vitro* and *in vivo* whole-cell recordings from PV interneurons showed that light dependent currents remained stable for periods of several seconds of illumination (Chow et al., 2010) (Fig. S1).

### Effect of silencing PV interneurons comprised of two phases

Photo-suppression of PV-mediated inhibition induced a significant (paired t-test, p<0.001, n=19 neurons) broadening of the tone frequency tuning curves (tuning width at half-maximum; Fig. 2a, b). Rescaling the tuning curve under PVoff conditions to the peak of the control tuning curve (Fig. 2a, green shading) highlighted the change in selectivity produced by silencing PV interneurons. These initial analyses suggested that tone-evoked PV inhibition contributes to sharpening response selectivity.

**Figure 2.**
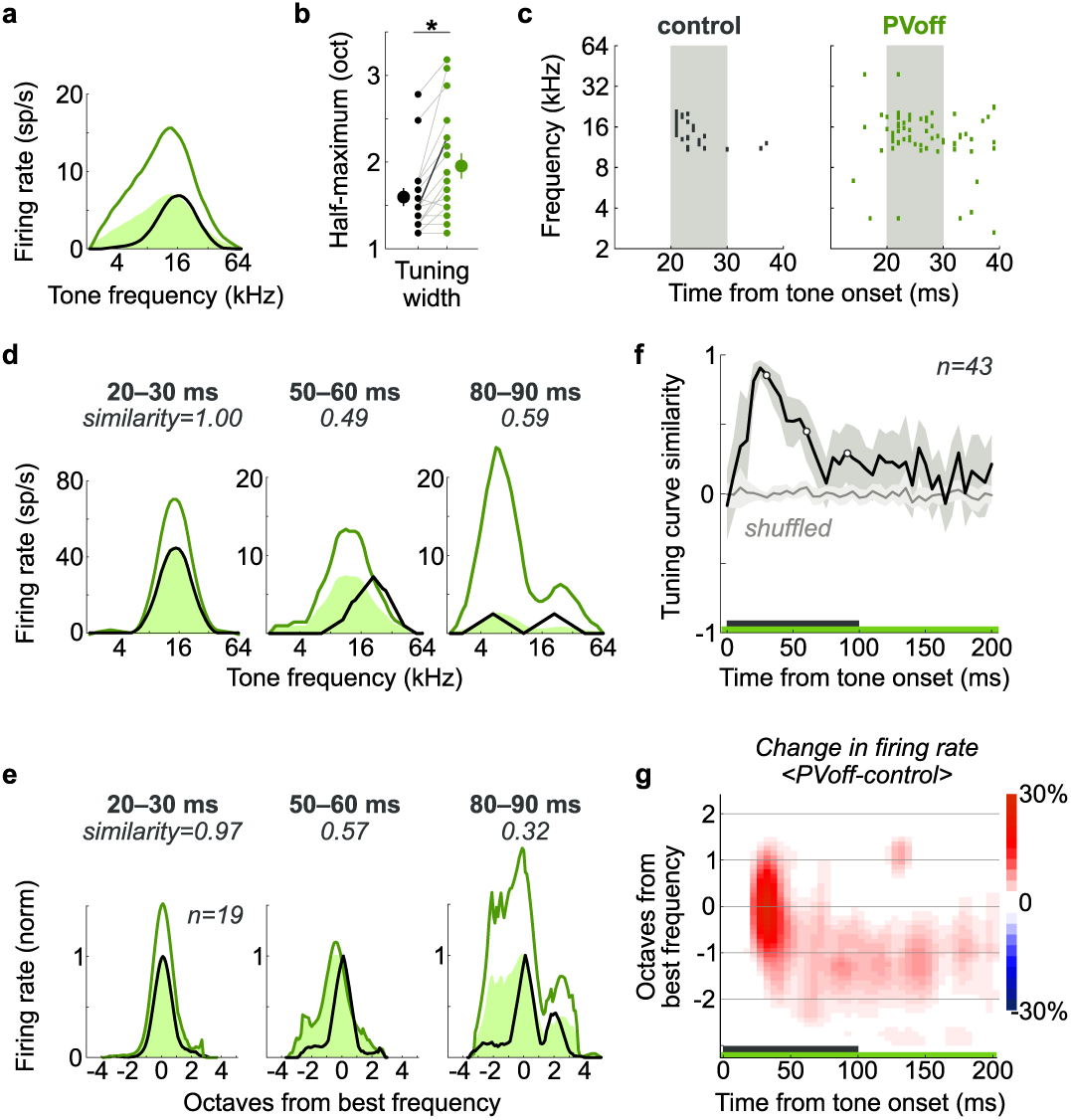
PV interneurons control the gain of the early component of tone responses. **a, b.** Photo-suppression of PV interneurons increased tuning width. Frequency response curves for the neuron in Fig. 1C were computed during the entire tone presentation (100 ms, a) under control {black) and PVoff (dark green) conditions. Tuning width computed at half maximum increased significantly across the population (b, n=19 neurons, p<0.001, paired t­ test). Black line in b corresponds to the tuning curves shown in a. **c.** Photo-suppression of PV interneurons increased the number of spontaneous and tone­ evoked spikes. Spike rasters for a single neuron show responses under control and PVoff conditions to 60 and 70 dB tones with frequencies ranging from 2 kHz to 64 kHz. Black bars indicate tone presentation; green bar indicates light presentation. Grey vertical shading shows 10ms window used for computing frequency response curves ind, e (20-30 ms). **d, e.** Photo-suppression of PV interneurons increased firing rate but did not change spectral selectivity of the early phase of tone-evoked responses. Frequency response curves for the neuron in a and average tuning curves for the population (e, n=19 neurons) were computed during 1Oms time windows starting at 20, 50, and 80 ms after tone onset under control (black) and PVoff (dark green) conditions. PVoff curves rescaled to the maximum of the corresponding control curve (light green) indicate changing similarity in spectral selectivity during the three 10 ms time windows. All frequency response curves were filtered with a one octave wide moving average filter. **f.** Across the population, photo-suppression of PV interneurons did not change spectral selectivity during the early phase of tone-evoked responses. Similarity between frequency response curves under control and PVoff conditions was computed in 1Oms time windows sliding in 5ms increments {black). Cosine similarity was calculated for independently normalized and mean-subtracted tuning curves obtained during each time window. For the shuffled control (grey), tone frequencies for each pair of tuning curves were independently shuffled before computing the cosine similarity. Shaded areas show s.e.m. Black bar indicates tone presentation; green bar indicates light presentation. **g.** During the later phase of tone-evoked responses, photo-suppression of PV interneurons increased neuronal activity predominantly away from the best frequency. Differences between normalized control and PVoff tuning curves were computed in 1Oms long time windows sliding in 5ms increments, then aligned by each neuron’s best frequency, and averaged. Tone-evoked activity started to increase (red) one to two octaves away from the best frequency approximately 50 ms after tone onset. Horizontal grey lines indicate one­ octave wide frequency bands. Black bar indicates tone presentation; green bar indicates light presentation.

Further analysis, however, revealed that the changes in frequency tuning caused by silencing PV-mediated inhibition evolved according to a characteristic temporal sequence. During the initial phase of the response (*20-30 ms*, Fig. 2d, e), the increase in evoked activity was not accompanied by a change in shape of the tuning curve. During this initial period, the tuning curve obtained in the PVoff condition could be rescaled to yield a precise fit to the control tuning curve (Fig. 2e, green shading), and tones that in the control condition failed to elicit a response also failed to elicit a response in the PVoff condition (Fig. 2c). For this example, the rescaling multiplier was 1.58, so that a maximum spike rate of 45 sp/s in the control condition became 71 sp/s in the PVoff condition. Thus, in the initial tone response PV interneurons perform pure gain modulation.

Later (>50 ms) in the response period, surprisingly, suppressing PV interneuron activity increased both the gain and width of the tuning curve. During this later phase the PVoff tuning curve could not be rescaled to fit the control tuning curve, and tones that in the control condition failed to elicit a response often elicited a response in the PVoff condition. Analysis of the temporal evolution of the tuning curves across the population confirmed that silencing activity of PV interneurons disrupted the shape of the tuning curves only about 50 ms after stimulus onset (Fig. 2f, g). Thus suppression of PV-mediated inhibition caused an early increase in the gain of the frequency tuning curve, followed by a late increase in both gain and width.

### Candidate synaptic mechanisms of reduced frequency selectivity

What is the mechanism by which PV-mediated inhibition exerts differential effects on the early and late components of the sensory evoked response? We consider two possible models. Both models posit that disruption of PV-mediated inhibition initially changes only the gain of the response. However, the additional late spikes elicited in the PVoff condition by tones that failed to elicit spikes in the control condition could arise from either a late decrease in inhibition, or from a late increase in excitation. According to the “late-inhibition” model, PV-mediated inhibition is normally recruited late in the sound-evoked response, and elimination of this late inhibition leads directly to changes in response selectivity during the late component. Alternatively, according to the “late-excitation” model, loss of early PV-mediated inhibition leads to inappropriate recruitment of excitation late in the response.

To distinguish between these models we used whole cell patch recordings to examine directly the changes in the time course of excitatory and inhibitory conductances caused by suppressing PV-mediated inhibition. We reasoned that if, as posited by the “late-inhibition” model, the late increase in response width is due to a reduction in a later component of PV-mediated inhibition, then it should be apparent as a reduction in the late component of the inhibitory conductance. If instead, as posited by the “late-excitation” model, the late increase in response width is due to a late increase in excitatory drive, then there should instead be an increase in the late component of the excitatory conductance with little or no change in the late inhibition.

### PV interneurons mediate inhibition evoked by thalamic stimulation *in vitro*

To establish the role of PV interneurons in providing thalamically driven feedforward inhibition in auditory cortex, we performed whole cell recordings in auditory thalamocortical slices (Cruikshank et al., 2002)(Fig. 3). Under control conditions, thalamic activation triggered overlapping inward and outward currents in excitatory neurons in thalamorecipient layers of auditory cortex (Cruikshank et al., 2002; Huang & Winer, 2000; Lee & Sherman, 2008; Muller-Preuss & Mitzdorf, 1984) (Fig. 3c). Decomposition of these overlapping currents into their underlying excitatory and inhibitory components (Wehr & Zador, 2003) revealed a stereotyped sequence consisting of a barrage of excitation followed quickly (1.5±1.3 ms) by inhibition, resembling responses to transient sensory stimuli in auditory cortex (Tan et al., 2004; Wehr & Zador, 2003; Zhang et al., 2003).

**Figure 3.**
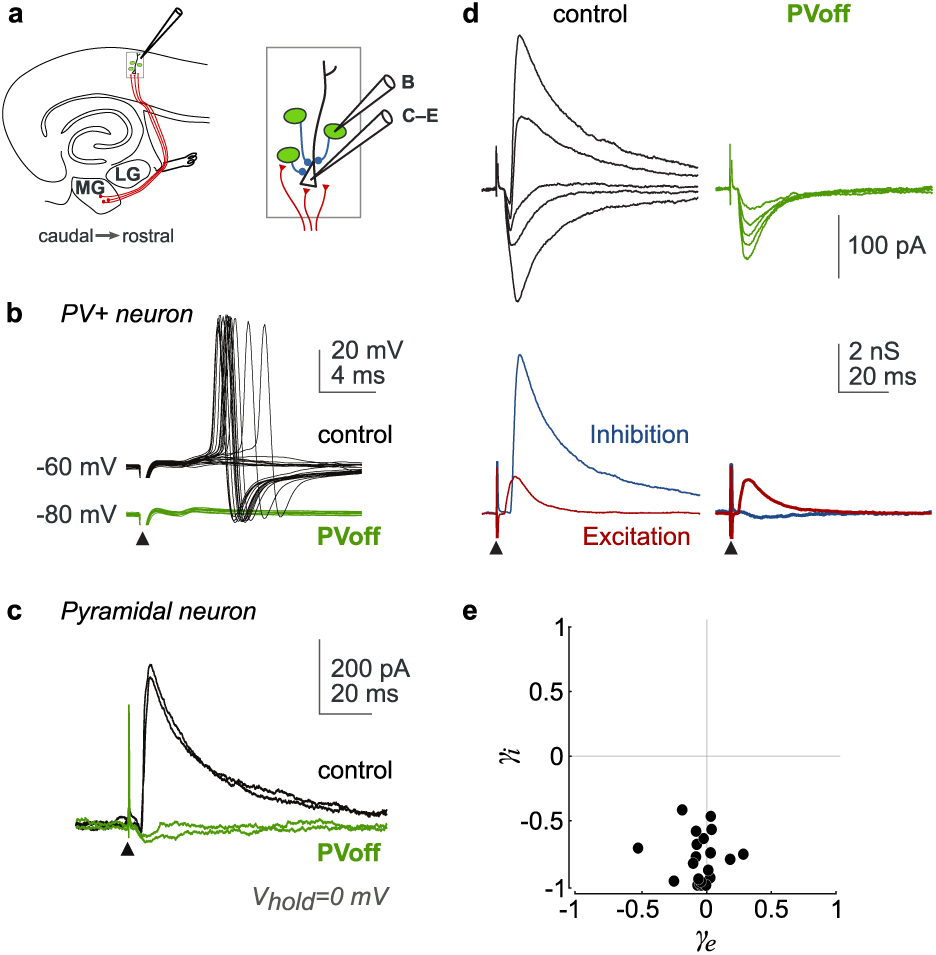
Silencing PV interneurons eliminated inhibition recruited by electrical activation of thalamocortical axons in vitro. **a.** Auditory thalamocortical slice preparation showing thalamocortical axons (red lines), stimulating electrode position, and whole cell patch recording configuration. Inset illustrates PV interneurons (green) making inhibitory synapses (blue) onto an excitatory pyramidal neuron. MG, medial geniculate and LG, lateral geniculate nuclei. **b.** Electrical activation of thalamocortical axons (triangle, 250 µA, 100 µs pulses) elicited spikes in a PV interneuron under control conditions (black). In interleaved trials, activation of Arch hyperpolarized the neuron by 20 mV and prevented spiking (PVoff, green) in response to identical stimuli. **c.** Strong outward synaptic currents elicited by thalamocortical stimulation under control conditions (black) upon thalamocortical stimulation (triangle, 200 µA, 100 µs pulses) were suppressed upon identical stimulation during continuous photo-activation of Arch (PVoff, green). Single trials were recorded from an Arch-negative pyramidal neuron (∼420 µm from the pial surface) held at the reversal potential for excitation. Spikes were blocked intracellularly with QX-314. **d.** Photo-suppression of PV interneurons (PVoff) blocked inhibitory (blue) but not excitatory (red) synaptic conductances evoked in a pyramidal cell by thalamocortical stimulation (triangle, 300 µA, 100 µs pulses). Synaptic currents (top, average of 6-8 repeats at five holding potentials) were recorded under control and test (PVoff) conditions, and further decomposed into excitatory (red) and inhibitory (blue) conductances (bottom). Spikes were blocked intracellularly with QX-314. **e.** Across the population (n=21 neurons), inhibitory change indices *(y;)* reflected suppression of inhibitory conductances *(y;=-0.80* +/- 0.18), while excitatory change indices *(Ye)* reflected little to no change in excitatory conductances (ye=-0.05 +/- 0.16). For the example ind, *y;=-0.97,* Ye=-0.05.

We then paired thalamic stimulation with photo-suppression of Arch-expressing PV interneurons (green, Fig. 3d). In this example, as in most neurons, silencing of PV-mediated inputs dramatically reduced the stimulus-evoked inhibitory conductance (blue, Fig. 3d) while the excitatory conductance remained largely unaffected (red, Fig. 3d). We quantified the change in inhibitory conductance as γi = (gi_PVoff_-gi_control_)/(gi_PVoff_+gi_control_), where g represents the peak evoked conductance, and defined the change in excitatory conductance (γe) similarly (see Methods). In most neurons (17/21) tested *in vitro*, inhibition was reduced by more than 75% during PV photo-suppression (γ_i_<-0.6), and in nearly half of the recorded neurons (10/21), inhibition was reduced by 90% or more (γ_i_<-0.82, Fig. 3e). Residual inhibition seen in some neurons in the PVoff condition could arise from technical factors such as incomplete photo-suppression or viral infection, or could represent a component of feedforward inhibition mediated by inhibitory neurons not expressing PV, as has been reported in somatosensory cortex (Porter et al., 2001; Tan et al., 2008). Nevertheless, these experiments demonstrate that PV interneurons are the dominant source of feedforward inhibition recruited in the auditory cortex by electrical activation of thalamocortical axons *in vitro*.

### Silencing PV interneurons reduces early tone-evoked inhibition

To demonstrate their action on postsynaptic neurons *in vivo*, we activated PV interneurons with Channelrhodopsin2 (see Methods) and performed voltage clamp whole-cell patch recordings. We decomposed the measured synaptic currents into excitatory and inhibitory conductances. These experiments confirmed that the action of PV-expressing neurons *in vivo* is inhibitory with a reversal potential close to calculated values (Fig 4b,c).

**Figure 4.**
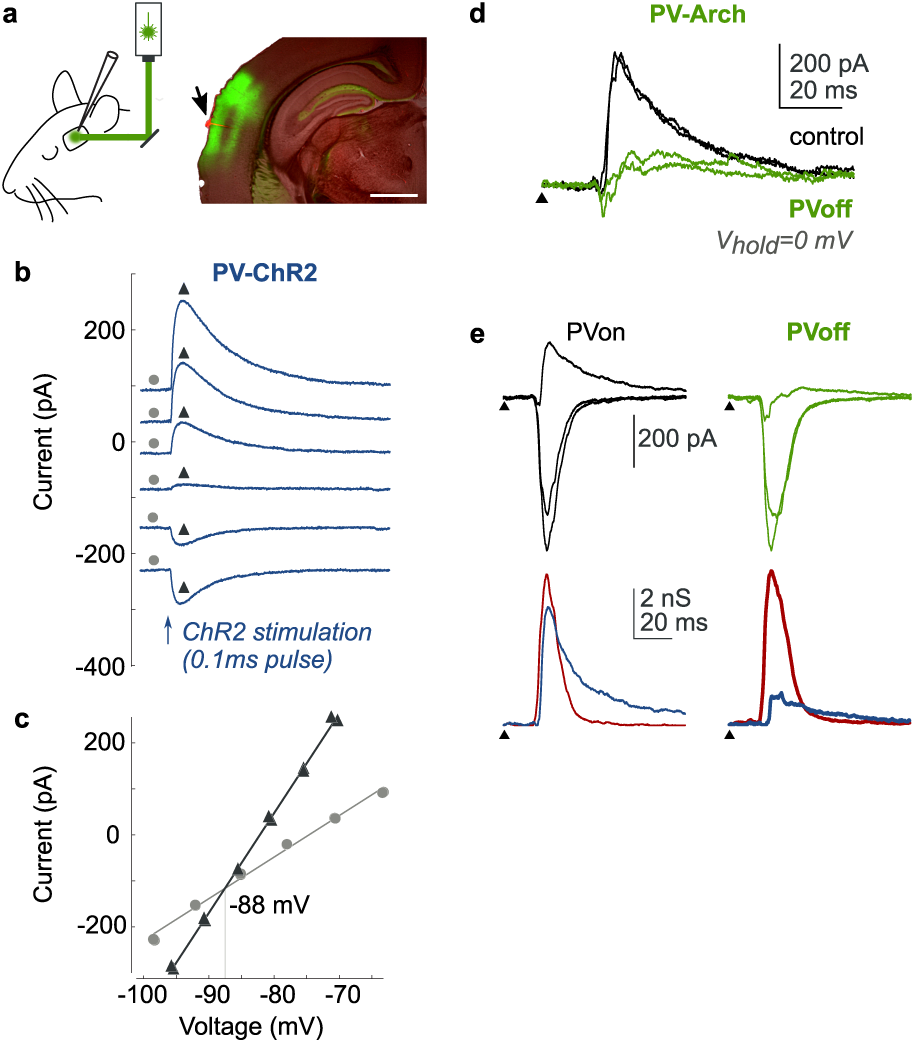
Silencing PV interneurons in vivo suppressed fast onset tone-evoked inhibition. **a.** In vivo preparation showing configuration of laser light stimulation and recording electrode (left). Blind whole cell patch recordings (arrow, red Di-I electrode track) were performed within the area of Arch-GFP expression (right, green area). Scale bar indicates 1 mm. **b.** Synaptic currents elicited by the brief ChR2-assisted activation of PV interneurons (0.1 ms, blue arrow) were purely inhibitory, showing reversal close to estimated values. **c.** Current-voltage relationship corresponding to traces in b. measured at times near baseline (circles) and peak inhibitory current (triangles). **d.** Outward synaptic currents (control, black) evoked by a tone (16 kHz, 70 dB SPL, 100 ms) were suppressed upon photo-suppression of Arch-expressing PV interneurons (PVoff, green). Single trials were recorded from an Arch-negative, presumed excitatory neuron (∼420 µm from the pial surface) held at the reversal potential for excitation. Because there is no current attributed to excitation at this potential, the current measured is directly proportional to inhibitory conductance. Triangle shows position of tone onset; light onset preceded tone by 250 ms. Spikes were blocked with QX-314. **e.** Photo-suppression of PV interneurons (PVoff) suppressed inhibitory (blue) tone-evoked synaptic conductances (tone onset at triangle) in a single neuron (same as ind). Synaptic currents (top, average of eight repeats) were recorded under control and test (PVoff) conditions, and further decomposed into excitatory (red) and inhibitory (blue) conductances (bottom).

Having established that strong inhibitory currents are imposed by PV interneurons on their cortical partners *in vivo*, we next returned to silencing PV interneurons while presenting tones. We used the blind whole-cell patch technique to uncover the underlying synaptic events evoked by the stimuli (Fig. 4d, e). As previously observed (Tan et al., 2004; Wehr & Zador, 2003; Zhang et al., 2003), pure tones (100 ms, 70 dB) elicited a rapid stereotyped sequence of excitatory and inhibitory conductances (Fig 4d,e). In interleaved trials we presented tones under control conditions and during photo-suppression of PV interneurons.

Under control conditions, excitation and inhibition were both elicited rapidly after the onset of the tone, with inhibition following excitation by a few milliseconds (Fig 4d, e, Fig 5a, b). Photo-suppression of PV-mediated inhibition caused a marked reduction in the early tone-evoked inhibitory conductance, consistent with the suppression of inhibitory interneurons. This reduction was maximal near the peak but persisted through the duration of the inhibitory conductance. In addition, after a delay of approximately 25 ms from tone onset, the excitatory conductance began to diverge from the control condition in the positive direction, becoming a statistically significant increase across the population for times between 30 and 200 ms after tone onset (Fig 5b,c). Thus PV photo-suppression resulted in a rapid reduction in inhibition followed by a more gradual increase in excitation.

**Figure 5.**
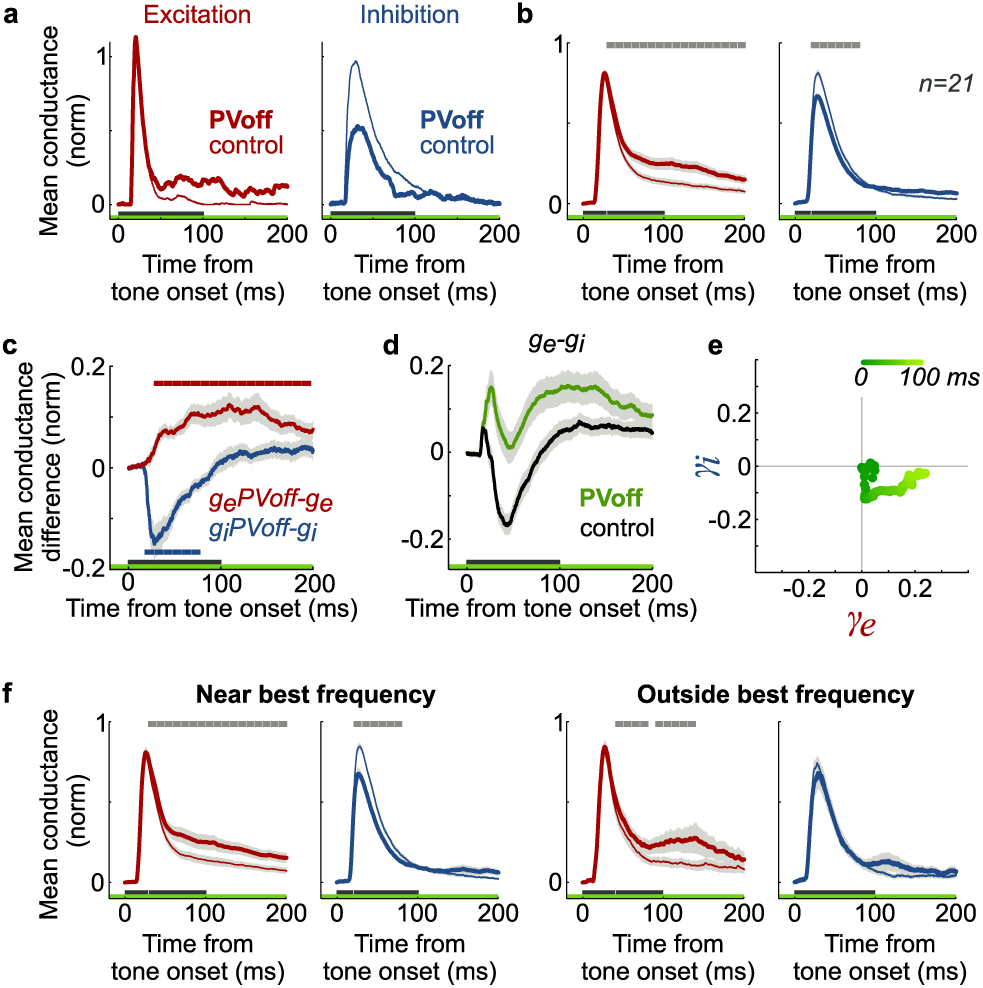
Silencing PV interneurons caused a late increase in tone-evoked excitation without decreasing inhibition for non-optimal frequencies. **a, b.** Excitatory conductances increased during the late phase of tone-evoked responses, while inhibitory conductances decreased during the early phase of responses. Excitatory (red) and inhibitory {blue) conductances in a single neuron (a) and across the population (b, n=21 neurons) recorded under control (thin) and PVoff {thick) conditions were aligned by tone onset, normalized to their corresponding control amplitudes and averaged. Green bars indicate light presentation, black bars indicate tone presentation, shaded areas show s.e.m. Shaded areas on top in b indicate significant difference between control and PVoff conditions evaluated in 10ms non-overlapping time windows (p<0.001, paired Wilcoxon signed rank test). **c.** An abrupt decrease of inhibitory (blue) conductances during the early part of tone presentation was followed by a gradual increase in excitatory (red) conductances across the population (n=21 neurons). Green bar indicates light presentation, black bars indicate tone presentation, shaded areas show s.e.m. Colored areas on top and bottom indicate significant change in excitation (red) or inhibition {blue) evaluated in 10 ms non-overlapping time windows (p<0.001, Wilcoxon signed rank test). **d.** Average synaptic ’drive’, estimated as difference between excitation and inhibition (ge-gi), was greater under PVoff condition (green) during the response. The early sharp peak corresponds to decrease in inhibition during the “window of opportunity”, whereas the late gradual peak corresponds to later increase in excitation (c). Green bar indicates light presentation, black bar indicates tone presentation, shaded areas show s.e.m. **e.** Average excitatory and inhibitory conductance change indices changed according to characteristic temporal sequence during the tone duration (100 ms). Decrease in inhibition dominated early during the response {dark green, *y;<O),* in contrast to increase in excitation late during the response (light green, *Ye>O)*. **f.** Left: Excitatory conductances (red) increased and inhibitory conductances (blue) decreased for tones near the best frequency for each neuron (n=21). Right: For tones more than half­ octave away from best frequency, excitatory conductances (red) increased, whereas inhibitory conductances {left) did not change. Green bars indicate light presentation, black bars indicate tone presentation, shaded areas show s.e.m. Shaded areas on top indicate significant differences between control and PVoff conditions evaluated in 1Oms non-overlapping time windows (p<0.001, paired Wilcoxon signed rank test).

We did not observe reliable differences between the responses of neurons recorded at different cortical depths, so we collapsed the data recorded *in vivo* across cortical layers (Linden et al., 2003). Since the majority of our recordings were performed above 550 μm, most of the neurons were likely recipients of direct thalamic input (Lee & Sherman, 2008).

The significant light-induced increase in firing rates and decrease in inhibitory conductances is consistent with effective optogenetic silencing of PV interneurons *in vivo*. The remaining portion of early tone-evoked inhibition could be provided by a different class of interneurons or by PV interneurons that were not effectively silenced. Factors likely contributing to incomplete silencing of PV interneurons include imperfect viral infectivity rates, strong but insufficient hyperpolarizing currents generated by Arch, and loss of light power at deeper cortical layers (Aravanis et al., 2007; Bernstein et al., 2008; Yizhar et al., 2011).

### Excitation increases late at non-optimal tone frequencies

The observed delayed increase in excitatory conductance supports the “late-excitation” model, according to which the late increase in the width of the tuning curve is an indirect consequence of altering the initial response gain. However, the persistent albeit smaller reduction in inhibition beyond the early phase means that these observations cannot rule out the “late-inhibition” model, which holds that the reduction of late component of inhibition is responsible for the loss of selectivity late in the response. To further distinguish between these two models we compared conductance changes near the best frequency (one-octave-wide band centered on the best frequency) and outside the best frequency (more than half-octave away from the best frequency) (Fig. 5f). Near the best frequency (Fig. 5f, left), the average conductance changes elicited by PV photo-suppression mirrored those of the entire population. However, outside the best frequency photo-suppression elicited only a late increase in excitation, without a significant change in average inhibitory conductance (Fig. 5f, right). Our data thus support the “late-excitation” model: the extra spikes elicited far from the best frequency late in the response arise from additional excitation, presumably unleashed by the loss of the normal precise regulation of the early response gain.

## Discussion

In the auditory cortex, the onset of a tone evokes strong excitation and inhibition that interact dynamically in time to shape frequency selective spike responses. We directly tested the role of early tone-evoked inhibition by manipulating the activity of a specific population of cortical interneurons, defined by the expression of PV. We demonstrated that this class of interneurons mediates early inhibition and then examined the role early inhibition plays in determining frequency tuning. We found that through enacting transient gain modulation, early inhibition controls long-term cortical dynamics, preventing late excitatory drive at non-optimal frequencies.

### PV interneurons mediate early stimulus-evoked inhibition

We hypothesized that manipulating the activity of PV interneurons would be an effective way to modify early tone-evoked inhibition. Many diverse interneuron classes coexist in the cortex and may serve different functions in regulating activity and processing sensory information (Markram et al., 2004; Moore et al., 2010). These interneuron types exhibit many differences including density of connections, regions targeted on the postsynaptic cell, strength of connectivity, and time of activation relative to the stimulus (Kawaguchi & Kondo, 2002; Kawaguchi & Kubota, 1997; Porter et al., 2001; Prieto et al., 1994; Tan et al., 2008). Of the different types of interneurons, PV-expressing interneurons emerge as ideal candidates to mediate early stimulus-evoked inhibition in many cortical areas, including auditory cortex (Celio, 1986; Cruikshank et al., 2007; Gibson et al., 1999; Hattori et al., 2017; Inoue & Imoto, 2006; Kerlin et al., 2010; Porter et al., 2001; Runyan et al., 2010; Schiff & Reyes, 2012; Sun et al., 2006; Wood et al., 2017). In the auditory cortex PV interneurons have high connectivity probabilities with nearby neurons, typically within the same lamina (Levy & Reyes, 2012; Yuan et al., 2011). Frequency selectivity is tonotopically arranged in the auditory cortex, thus PV interneurons likely inhibit neurons sharing similar frequency preferences (Bandyopadhyay et al., 2010; Katzel et al., 2011; Yoshimura & Callaway, 2005; Yuan et al., 2011). PV interneurons may have a wider tuning bandwidth than excitatory neurons (Atencio & Schreiner, 2008; Wu et al., 2008); c.f. (Moore & Wehr, 2013), but it is unclear whether this difference plays a functional role. We directly demonstrate that selective silencing of PV interneurons eliminates significant portions of the inhibitory conductance experienced by an auditory cortical neuron after the activation of the auditory thalamus, either by direct electrical stimulation *in vitro* or by the onset of a tone *in vivo*.

### Role of inhibition in initial frequency tuning

Response fields in many cortical areas widen after topical application of pharmacological inhibitory blockers (Dykes et al., 1984; Sillito et al., 1980; Wang et al., 2000; Wang et al., 2002) or after optogenetic suppression of inhibition (Aizenberg et al., 2015), suggesting that inhibition sharpens the selectivity of responses. Inhibition is then assumed to be tuned more widely than excitation and to suppress responses to non-preferred stimuli (Calford & Semple, 1995; Lee et al., 2012; Liu et al., 2011; Swadlow & Gusev, 2000). However, intracellular studies showing that excitation and inhibition are largely co-tuned predict that inhibition does not play a major role in shaping response selectivity (Anderson et al., 2000; Atallah et al., 2012; Borg-Graham et al., 1998; Tan & Wehr, 2009; Wehr & Zador, 2003; Wilson et al., 2012; Zhang et al., 2003; Zhou et al., 2014).

We find direct evidence that in the auditory cortex the frequency tuning of the initial excitatory barrage determines the frequency tuning of the recipient neuron. Tuning curves broadened late in the tone response as an indirect consequence of blocking early inhibition. In addition, we did not observe changes in first spike latency upon suppression of early inhibition, further supporting major role of excitation in the initial frequency tuning.

### Temporal dynamics of auditory responses

In some respects, our results are similar to findings in visual cortex, which show that PV interneurons modulate the response amplitude of neurons in mouse visual cortex (Atallah et al., 2012; Lee et al., 2012; Wilson et al., 2012). But the role of PV interneurons in modifying more complex response properties, such as orientation tuning, remains controversial. Some studies suggest that PV interneurons do not affect orientation tuning in V1 (Atallah et al., 2012; Nelson et al., 1994; Wilson et al., 2012), while others report changes in orientation tuning mediated by PV interneuron activity (Lee et al., 2012).

An important difference between the auditory cortex and the visual cortex is the temporal dynamics of both stimuli and responses (King & Nelken, 2009). Experiments in the visual cortex typically involve measuring spike rates over much longer time periods, on the order of seconds. Such measurements are appropriate for areas where neurons readily fire in sustained fashion in response to prolonged sensory stimuli. In contrast, auditory stimuli can be very brief, contain fast temporal modulations, and involve quickly changing statistics. In turn, auditory responses are often transient under control conditions.

If the tone responses were measured throughout the tone and without direct conductance evidence to the contrary, the tuning width increases that we observed might be interpreted as evidence for lateral inhibition at non-preferred frequencies. However, the tuning width changes occur tens of milliseconds after the initial “window of opportunity” and are associated with increased excitation rather than decreased inhibition. Previous predictions could only address the early effects of inhibition during the initial response (Wehr & Zador, 2003; Zhang et al., 2003). Here we show that transient early changes can affect activity in a specific manner at much later time points.

Our results indicate that gain modulation can have more far reaching effects than simply imposing a linear shift in firing rates. By quenching the response in time, early inhibition from PV interneurons prevents late accumulation of excitation at non-optimal frequencies. This tight temporal control might also enhance the response fidelity to more complex, temporally modulated sounds. Together, our data suggest that the proper functioning of auditory cortical circuitry depends on the temporal dynamics of stimulus-evoked responses, and that transient alterations in activity can have long lasting consequences.

## Methods

All experiments were performed in strict accordance with the National Institutes of Health guidelines, as approved by the Cold Spring Harbor Laboratory Institutional Animal Care and Use Committee.

### Virus production

To prepare the construct for producing the adeno-associated virus encoding cre-dependent Archaerhodopsin-3 (Arch), we used an AAV backbone (from Dr. Karl Deisseroth, Stanford) containing the DNA sequences of the EF1α promoter, FLEX-configured loxP and lox2722 sites (Atasoy et al., 2008), and woodchuck post-transcriptional element (WPRE) (Loeb et al., 1999). DNA encoding ss-Arch-eGFP-ER2 (from Dr. Edward Boyden, MIT) consisted of a membrane targeting signaling sequence (ss) and an endoplasmic reticulum export signal (ER2) joined to a sequence coding for Arch fused to enhanced green fluorescent protein (eGFP) (Chow et al., 2010). This sequence was amplified using standard PCR methods and ligated into the AAV backbone. The resulting construct (pAAV-EF1α-FLEX-ss-Arch-eGFP-ER2-WPRE) was sent to the Joint Vector Laboratories (The University of North Carolina at Chapel Hill Gene Therapy Center) for production of high titer AAV (serotype 2/9).

For experiments involving the expression of cre-dependent Channelrhodopsin2 (ChR2) (Fig. 4) an AAV construct containing ChR2 in the FLEX configuration (pAAV-EF1α-FLEX-ChR2-eYFP-WPRE) from the lab of Dr. Karl Deisseroth (Stanford) was amplified and sent to the Penn Vector Core (University of Pennsylvania Gene Therapy Program) for production of high titer AAV (serotype 2/9).

### Directing Arch expression

For viral injection experiments, young (age P15–25) PV-IRES-cre mice (stock number 008069, The Jackson Laboratory) (Hippenmeyer et al., 2005) were injected with virus (Lima et al., 2009) in 5–8 small craniotomies made along the lateral extent of the dorsal surface of the skull aligned with the rostro-caudal location of left auditory cortex. For a single track of virus injection, infected PV interneurons were typically limited to within 500 μm from the center of the track. Many individual injection tracks were thus required to span the region of auditory cortex. Virus was delivered by pressure ejection (5–10 ms pulses at 20–25 psi, controlled by Picospritzer II, General Valve, Fairfield, NJ) from a glass pipette (∼15 μm tip diameter) lowered to the vertical depth of the auditory cortex. The craniotomies were typically filled with silicone sealant (Kwik-Cast, World Precision Instruments) and the skin closed with liquid suture (VetBond, 3M). Animals were removed from the anesthesia, treated with ketofen for analgesia (5 mg/kg), and returned to their home cages after regaining movement.

For a subset of *in vitro* experiments we used transgenic mice expressing Arch in PV interneurons. To generate these mice we bred PV-IRES-cre mice with Ai35D mice (Dr. Hongkui Zeng, Allen Institute for Brain Science), in which the CAG promoter, a lox-stop-lox cassette, and the ss-Arch-GFP-ER2 sequence are inserted into the Rosa locus (Madisen et al., 2012). When necessary, we genotyped offspring for a portion of the Arch sequence with custom designed PCR primers (fwd: CCATCGCTCTGCAGGCTGGTT, rev: CAAGACCAGAGCTGTCAGGGTGT TA).

### Acute slice preparation

Auditory thalamocortical slices (Cruikshank et al., 2002) were obtained from mice older than postnatal day 35. Mice were deeply anesthetized with a mixture of ketamine (120 mg/kg) and medetomidine (0.5 mg/kg) and transcardially perfused with chilled artificial cerebrospinal fluid (ACSF) containing (in mM) 127 NaCl, 25 NaHCO_3_, 25 d-glucose, 2.5 KCl, 1 MgCl_2_, 2 CaCl_2_ and 1.25 NaH_2_PO_4_, aerated with 95% O_2_ 5% CO_2_. Brains were extracted and quickly submerged in chilled cutting solution containing (in mM) 110 choline chloride, 25 NaHCO_3_, 25 d-glucose, 11.6 sodium ascorbate, 7 MgCl_2_, 3.1 sodium pyruvate, 2.5 KCl, 1.25 NaH_2_PO_4_ and 0.5 CaCl_2_, aerated with 95% O_2_ 5% CO_2_. Slices 500 μm thick were prepared using a vibratome in a chamber cooled to 2°C (Microm, Walldorf, Germany) and transferred into a holding chamber circulating warmed ACSF. The slices were incubated in the holding chamber at 32–34 °C for 30–45 min and then held at room temperature (22–25 °C) for the remainder of the experiment. Electrophysiological recordings were performed in a submersion style recording chamber with circulating ACSF at 28-30 °C. Neurons were visualized for patch recordings using standard infrared differential interference microscopy (Olympus, Center Valley, PA) and a CCD camera (Optronics, Goleta, California).

### Anesthetized in vivo preparation

Ten to fourteen days after viral injection, PV-IRES-cre mice were anesthetized with isoflurane (2% isoflurane for induction, 0.5–0.75% for maintenance of anesthesia), transferred to a custom naso-orbital restraint and placed on a heating pad during the recording session. A cisternal drain was performed and a craniotomy and durotomy were performed above the left auditory cortex. The area of the greatest GFP expression was determined visually under a fluorescence microscope and targeted for recordings. Standard blind *in vivo* whole-cell (n=27) and cell-attached (n=23) recordings were performed in all subpial depths (230–860 μm), as determined by micromanipulator travel; 75% of all recordings were <550 μm.

### Electrophysiological recordings

For both *in vitro* and *in vivo* experiments, patch electrodes were pulled from filamented, thin-walled, borosilicate glass (outer diameter, 1.5 mm; inner diameter, 1.17 mm; World Precision Instruments, Sarasota, FL) on a vertical two-stage puller (Narishige) and filled with internal solution containing (in mM): 140 KGluconate, 10 HEPES, 2 MgCl_2_, 0.05 CaCl_2_, 4 MgATP, 0.4 NaGTP, 10 Na_2_-Phosphocreatine, 10 BAPTA, and 6 QX-314, with pH adjusted to 7.25 and diluted to 290 mOsm. Resistance to bath was 3.5–5.0 MOhm before seal formation. Recordings were obtained using Axopatch 200B amplifiers (Molecular Devices, Sunnyvale, CA) set to lowpass filter at 2–5 kHz and a custom data acquisition system written in MATLAB (Mathworks), with a sampling rate of 10 kHz.

To estimate excitatory and inhibitory conductances three to seven holding potentials were used. At each potential, after a one second equilibration period, five to ten 10 mV voltage pulses were delivered to monitor series resistance and input resistance, followed by electrical or tone stimulation either alone or accompanied by light stimulation (see below).

### Light stimuli

For *in vitro* experiments, green light pulses were delivered via a conventional fluorescence illuminator (Xcite 120, Lumin Dynamics) passed through a 535/50 nm excitation filter (Chroma 4100) and focused onto the tissue through a 60x objective with a field diameter of 350 μm. The total power was approximately 30 mW. A shutter was used to control the temporal gating of the light delivery (Uniblitz, Rochester, NY).

For *in vivo* recordings, green light pulses were delivered using either a 30 or 50 mW, TTL gated 532 nm laser (1 mm beam width, Extreme Laser, China). For experiments during which ChR2 was photo-activated (Fig. S1), blue light pulses were delivered using a TTL gated 473 nm laser (1mm beam width, Lasermate Group). In both types of experiments, a small mirror on a manipulator was used to reflect the laser beam to the recording site, achieving an illuminated area of approximately 1 mm diameter centered at the tip of recording electrode. The illuminated area was thus large enough to cover the entire primary auditory cortex (Stiebler et al., 1997); (Linden et al., 2003).

### Electrical stimuli

Constant current pulses, 50–400 μA and 100 μs in duration, were delivered by a twisted bipolar microelectrode (0.5 MΩ, World Precision Instruments, Sarasota, FL) driven by an ISO-Flex stimulus isolation unit controlled by a Master-8 pulse generator (A.M.P.I., Jerusalem, Israel). In interleaved trials electrical stimuli were presented during light, where the onset of light preceded the electrical stimuli by 250 ms.

### Sound stimuli

All *in vivo* experiments were conducted in a double-walled sound booth (Industrial Acoustics Company). Free-field acoustic stimuli were presented at a 200 kHz sampling rate using a custom real-time Linux system driving a Lynx L22 audio card (Lynx Studio Technology Inc, Newport Beach, CA) connected to an amplifier (Stax SRM 313, STAX Ltd, Japan), which drove a calibrated electrostatic speaker (TDT Alachua, FL) or a custom-built speaker located 8 cm lateral to, and facing, the contralateral (right) ear.

Sound stimuli consisted of a fixed pseudo-random sequence of 100 ms long pure tones with 5ms 10–90 % cosine-squared ramp was presented with inter-tone interval of 2 s. For whole-cell recordings we used pure tones of 4–48 kHz (four per octave) presented at 70 dB SPL. For cell-attached recordings we used pure tones of 51 frequencies logarithmically spaced between 2–64 kHz (ten per octave) presented at 0–70 or 10–70 dB SPL (at 10dB steps). Every other tone was accompanied by a flash of green light, which started 250 ms before tone onset and ended 300 ms (whole-cell recordings) or 150 ms (cell-attached recordings) after tone offset.

### Synaptic conductance analysis

In both *in vitro* and *in vivo* recordings, series resistance was computed at each holding potential using the peak current transients evoked by five to ten -10 mV square voltage pulses. No online series resistance compensation was used. Holding potentials were corrected for a calculated liquid junction potential of 12 mV. Total synaptic conductance, corrected for series resistance, was then computed assuming an isopotential neuron, and further decomposed into excitatory and inhibitory conductances as previously described (Wehr & Zador, 2003).

Synaptic conductance responses were defined as the maximum conductance amplitudes of the excitatory or inhibitory conductance waveforms evoked by electrical or sound stimulus presentation. To compare responses under control conditions and during photo-suppression of PV interneurons (PVoff), we computed the inhibitory conductance change index (γi) as γi = (gi_PVoff_-gi_control_)/(gi_PVoff_+gi_control_) where g refers to the peak evoked conductance and subscripts indicate the control (with light off) and test (PVoff, with light on) conditions. The excitatory conductance change index (γe) was defined similarly. Excluded from this analysis were evoked conductances smaller than 0.5 nS in control conditions. For *in vivo* experiments, conductances were also required to reach half maximal amplitude within 10 ms to 50 ms after tone onset.

To compute population average of excitatory and inhibitory conductances across the populations of neurons (Fig. 5) we first normalized each pair of conductance waveforms to their control condition amplitudes and then averaged them. Included in this analysis were conductance waveforms with control condition amplitudes of at least 0.5 nS from neurons with γi<=-0.11 for at least one tone frequency (corresponding to 20% decrease in inhibitory conductance, n=21 out of 27 neurons). Statistical significance of difference between conductance waveforms was computed using Wilcoxon paired, two-tailed signed-rank test in non-overlapping 10ms windows. To correct for multiple comparisons we used p=0.001 as the threshold for significance.

### Spike response analysis

Spikes recorded in cell-attached mode were extracted from raw voltage traces by applying a high-pass filter and thresholding. Spike times were then assigned to the peaks of suprathreshold segments and rounded to the nearest millisecond. Spiking responses were defined as mean spike firing rate during a set time window. Tuning curves at a given sound intensity were computed by first computing mean spiking responses to each tone frequency, subtracting spontaneous rate from each response, and then smoothing the resulting tuning curve with zero-phase one-octave-wide (ten neighboring frequencies) moving average filter (filtfilt in Matlab). Only tuned tone-responsive neurons were included in the analysis (n=19 out of 23 neurons).

Population tuning curves were estimated by first computing, for each neuron, tuning curves for responses to 60 and 70dB tones under control and PVoff conditions. Best frequency for each neuron was then defined as the frequency corresponding to the peak of control tuning curve. Each pair of tuning curves was then normalized to the best frequency response under control condition, tuning curves were aligned by their best frequencies and averaged (Fig. 2e).

To compute tuning curve similarity (Fig. 2f), we first computed pairs of tuning curves as above (control, PVoff) from spikes in a sliding 10ms time window. In each time window, each tuning curve was normalized to its corresponding maximum value. Cosine similarity between the tuning curves was then computed as the cosine of the angle between the two mean-subtracted tuning curves (T_control_ and T_PVoff_) represented as vectors, i.e. similarity= (T_control._T_PVoff_)/(||T_control_||||T_PVoff_||), where (T_control._T_PVoff_) is the dot product and ||T_control_||, ||T_PVoff_|| are the Euclidean norms of T_control_ and T_PVoff_.

All error bars are s.e.m. unless noted otherwise.

## Acknowledgements

This work was supported by grants from the Slovak Research and Development Agency APVV-19-0585 to TH and a Swartz Postdoctoral Fellowship for Theoretical Neuroscience to APR.

