## Supplemental Figure 1 for "Early dynamics of excitation and inhibition maintain late frequency tuning in auditory cortex"

### Supplemental information

Ashlan P Reid[1], Anthony M Zador[2] and Tomáš Hromádka[3]
*[1]Department of Physiology, New York Medical College, Valhalla, NY 10595*
*[2]Cold Spring Harbor Laboratory, 1 Bungtown Road, Cold Spring Harbor, NY 11724*
*[3]Institute of Neuroimmunology, Slovak Academy of Sciences, Bratislava, Slovakia*

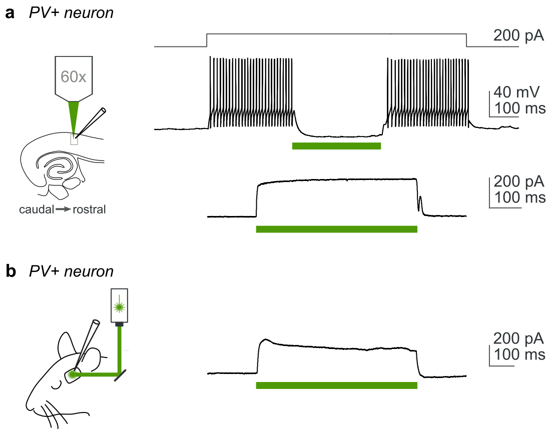


**Supplemental Figure 1**
Light dependent currents remained stable during illumination.

In vitro (**a.**) and in vivo (**b.**) recordings from PV interneurons showed that continuous light illumination suppressed firing (top) and light dependent currents remained stable for periods of hundreds of milliseconds of illumination (green bars)
